# Simulating Alpha Particle Doses at the Micron Scale from Prostate Cancer Patient Derived Bone Metastatic Biopsies Using GATE

**DOI:** 10.1101/2024.12.11.627943

**Authors:** Abdulaziz Said, Mahdjoub Hamdi, Isabella Salerno, Nadia Benabdallah, Norman F. Turtle, Diane Abou, M. Allan Thomas, Justin Mikell, Daniel Thorek

## Abstract

Radiopharmaceutical therapies are poised to enhance patient care for several currently untreatable metastatic cancers. Radium-223 dichloride citrate is indicated for treatment of bone metastatic castrate resistant prostate cancer, serving as the primary use of alpha particle emitting radium to irradiate bone lesions. Improvements and refinement of such therapies relies on patient centric and micro scale quantification to assess and compare efficacy. Computational modeling and Monte Carlo simulations provide a valuable tool for understanding micro scale phenomena of radiopharmaceutical therapies. Via simulation, we undertake and illuminate dose profiles of radium-223 dichloride treatment, evaluating energy and dose distributions based on primary, patient-derived specimens. A set of four activity distributions were simulated on three patient bone lesion biopsy samples. These simulations validate the novel tool for micron-scale modeling with patient-derived specimens. Ablative dose profiles are shown to be driven by uptake distributions as well as the target tissues’ microstructure.

## Introduction

Recent decades have seen the introduction of myriad new cancer treatments and therapy modalities. These innovations have led to a 33% decrease in cancer mortality driven in part by over 100 newly approved drugs by the United States Food and Drug Administration (FDA) between 2003 and 2021, with the greatest improvement in leukemia, melanoma, kidney cancer, and lung cancer (Michaeli, Michaeli, and Michaeli 2023). Ionizing radiation can be lethal to cancer cells, which forms the basis for this key pillar of cancer care. As such, radiation therapies have especially been foundational in curing patients of disease, contributing to 40% of curative treatments (Baskar et al. 2012).

Traditional radiation therapies (RT) either leverage external beams of high energy photons or locally implanted sealed source brachytherapy devices in order to irradiate tumors and avoid organs at risk. Although refinements in traditional RT have been transformative in treating and curing patients, it is a loco-regional treatment paradigm and remains effective primarily for cancer which has not yet spread through the patient. Metastatic disease, characterized by numerous lesions throughout the patient, remains largely uncurable and is the cause of significant mortality each year. External beam radiotherapies are sparingly used in the metastatic setting is due to efficacy and safety. First, micro metastases may not be visible on radiological scans and therefore are often missed. Further the high number of lesions spread throughout the body makes it difficult to avoid toxicity with external RT modalities due to entrance and exit absorbed doses and the invasive nature of sealed source brachytherapy.

Targeted delivery of radioactive isotopes to disease sites can enable more biologically precise and localized lesion irradiation. Thus, this approach may bridge the gap between effective radiation treatments and disseminated disease. This approach is known as targeted radiotherapy (TRT) and leverages the unique properties of targeted ligands coupled with beta or alpha particle emitting nuclides for these purposes. There is considerable interest in the use of high linear energy transfer alpha particle emissions. These alpha particles (helium nuclei) are released upon decay from heavy isotopes and deposit megaelectron volt (MeV) energies along a short path length in tissues, of approximately several cell-diameters (Brechbiel 2007). This provides the capability to deposit lethal radiation doses to a highly circumscribed region. Accomplishing this task requires the engineered molecule to both emit localized radiation and selectively target cancerous sites while avoiding significant uptake in normal tissue.

The first and only targeted alpha therapy approved for use to-date is Radium dichloride citrate (^223^RaCl_2_), which is used to treat bone metastatic castrate resistant prostate cancer (bmCRPC) (Kluetz et al. 2014). This aggressive late-stage form of prostate cancer is the most common noncutaneous cancer in males and a leading cause of cancer mortality (Siegel, Giaquinto, and Jemal 2024; Logothetis et al. 2018). Prostate cancer most frequently metastasizes to the bone (Lange and Vessella 1998; Sathiakumar et al. 2011). In this therapeutic strategy, Radium is known to be a bone-seeker, and it preferentially localizes to sites of active bone turnover irradiating bmCRPC lesions (Abou et al. 2016; Henriksen et al. 2002). Presently, dosing of Radium Dichloride treatment is determined by a patient’s weight and is set to 55 kBq/kg, administered intravenously in six cycles separated by four-week intervals (Sternberg et al. 2020; Humm et al. 2015). These are exceptionally low levels of administered activity, relative to activities administered for diagnostic procedures or other radiotherapies. This demonstrates the potency per unit of activity for Radium-223 relative to Lu177 PRRTs with prescribed fixed activities of 7.4 GBq per cycle (Lutathera, Pluvicto). Ra-223 emits four alpha particles through the decay chain depicting in Figure 1.

**Figure 1.**
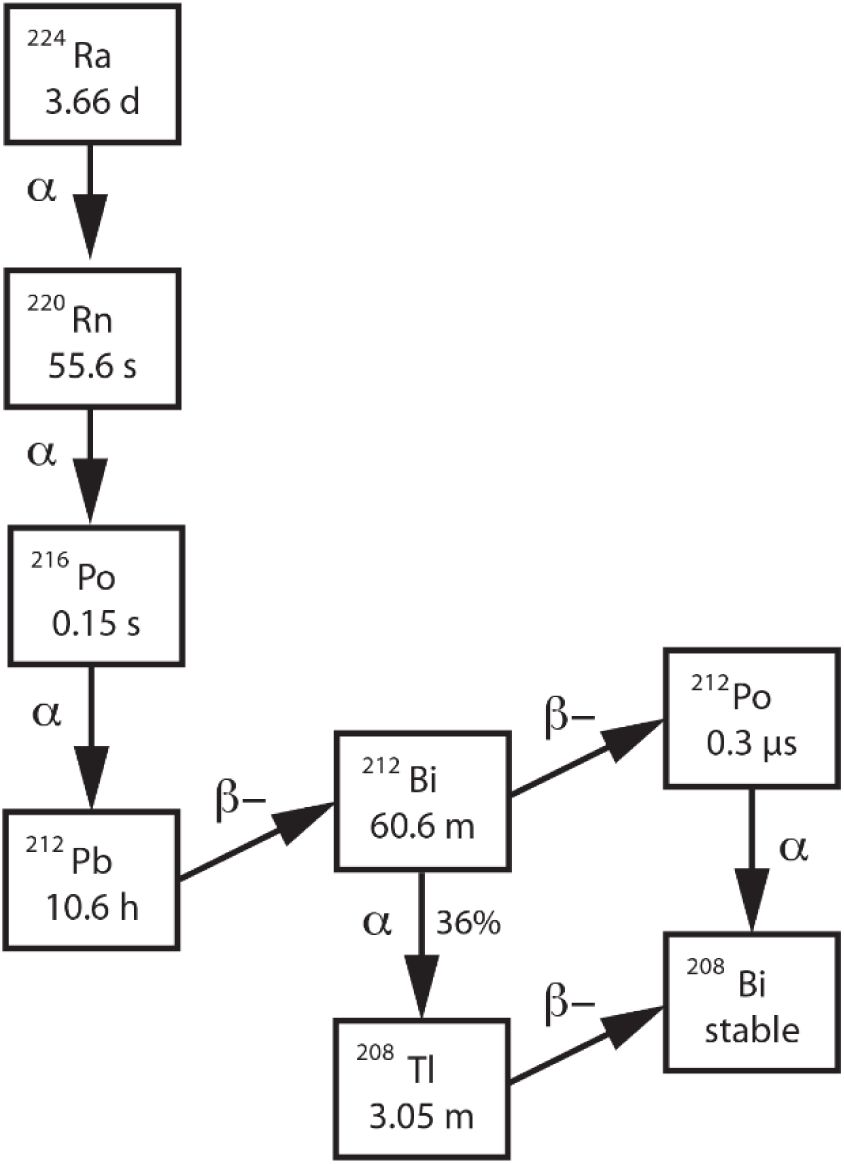
Decay Scheme of Radium 223 and Progeny

Despite the significant promise of 223Ra and other alpha particle emitting therapies, there remains much that is unknown about the localized absorbed doses and methods for optimization of this emerging drug class. Improvements in TRT development have relied on a mixture of *in vitro* and *in vivo* studies to establish drug characteristics, efficacy, and pharmacokinetics.

Traditional methods have focused on whole organ estimations of absorbed dose (Sgouros et al. 2010). This approach presents limitations as micro level heterogeneities are not considered. Due to alpha particles short penetration in tissue and the non-uniform uptake of such emitters at sites of disease, microscopic shifts in deposited energy can lead to significant irregularities in cancer cell death. Computational methods have aimed to quantify micro level interactions through *in silico* simulations of these therapies with Monte Carlo (MC) simulations garnering special attention (Hobbs et al. 2012; Pinto et al. 2020; Koniar et al. 2023).

Although MC techniques have been gaining in popularity for dosimetry calculations, there exists a paucity of information from preclinical animal studies regarding the critical input of radionuclide micro distribution, and even less from patient samples over preclinical sources.

Hence, the purpose of this work has been to perform microscale dosimetry estimates on patient biopsies. Here, we leverage a computerized method using high resolution autoradiography from patient bmCRPC biopsies. Bone architecture is delineated using high resolution micro-X-ray computed tomography (micro-CT) and high-resolution radionuclide distributions through digital autoradiography (DAR) (Benabdallah et al. 2024). Using the Geant4 simulation toolkit for particle transport through matter, extensively validated in biomedical applications (Agostinelli et al. 2003) and the Geant4 Application for Tomographic Emission, or GATE (Jan et al. 2004), 223Ra therapy has been simulated with special analysis on dosimetry measures. Using known total activity contained in the sample from well counting, we have distributed activity in different assumed spatial configurations reporting the subsequently simulated absorbed doses; furthering understanding of intra and inter-lesional in dosimetry.

## Materials and Methods

### Sample preparation

Six bmCRPC patients undergoing standard radium dichloride treatment were biopsied at bone lesion sites 24 hours after administration of fifty-five kBq/kg of Radium-223, under an Institutional Review Board approved protocol # 20141115. Patients received pretreatment 99mTc-methyl diphosphonate bone scans, and candidate osseous lesions were identified. CT-guided percutaneous drill-assisted biopsy samples using the coaxial OnControl system (Teleflex Arrow) were then extracted. The samples were embedded in optimal cutting temperature (OCT) embedding media, frozen and stored at -80 °C, fully described in our previous work (Benabdallah 2024).

Samples were then scanned at high resolution using a microscopic X-ray computed tomography (micro-CT) device, with temperature maintained using dry ice. Images of the excised biopsies were taken using a high-resolution micro-CT (VivaCT40; Scanco). At least two biopsy samples were extracted per patient and micro-CT images were collected for each sample at a 17.5-micron isotropic voxel resolution (70 kVp, 114 μA). Biopsy samples were γ-counted using an open-window protocol for 10 min using a National Institute of Standards and Technology source– calibrated system (Wizard2; Perkin Elmer). These processed CT volumes and the activity measured from the gamma counter were then used as input for simulations of Radium-223 treatment.

### Simulation preparation

The micro-CT images were optimized for use in simulations. First, each image was cropped, and all Hounsfield unit (HU) values were restricted to a max value of three thousand. Voxels were then categorized as either bone, bone surface, or soft tissue, according to consensus literature, namely the categorization of voxels with a value between 1,000 and 2,000 as bone, and while those with values under 1,000 were set as soft tissue (Richard Bib et al. 2015). The bone tissue interface was defined as a subset of the bone voxels, which is the voxels identified as bone and also adjacent to a soft tissue voxel. These processed CT images were used as input for simulation of Radium-223 treatment. The Geant4 Application for Tomographic Emission (GATE) is a widely used package for simulating the physics of nuclear processes through matter. GATE, an application built over the Geant4 software suite gives users the ability to simulate decay schemes over specified volumes and materials via user friendly command line macro files (Jan et al. 2004). In this investigation, GATE was employed to simulate the entire Radium-223 decay chain generating dose absorption maps which could then be rapidly visualized and analyzed.

Generation of GATE macro files has been automated via a Python application for each biopsy sample.

### Simulation architecture

We used the GATE ion source in order to simulate Radium-223 decay. The ion source allows for the simulation of any ion through the definition of its atomic number (Z), and atomic weight (A), set here to 88 and 223, respectively. This is the most realistic decay scheme as both radioactive decay and atomic de-excitation are simulated. The physics processes were controlled via use of the radioactive decay module, QGSP_BIC_EMZ and emstandard_opt4 physics lists, with use of the James Random Ranlux64 MersenneTwister engine. Each run was set to have a total activity of 25 Bq determined via estimation from previously NIST-calibrated gamma counting absolute quantification (Benabdallah et al. 2024), and a run time of 10 minutes as initial irradiation times. An additional set of simulations was run for one sample and 2.4 hours and compared with that of the ten-minute simulation. For each biopsy sample, a variety of source distributions were simulated. These included a point source with all activity confined to a single voxel; activity evenly distributed throughout the source region; and activity distributed unevenly. The site of activity uptake was set to be the bone/tissue interface. For GATE to simulate voxelized sources in this manner, two input files are required per simulation. A voxelized source file with voxels containing source activity labeled, and an activity parameter file to describe the amounts of activity set in each of the labeled voxels. These files were generated using an automated Python script with the biopsy micro-CT images as inputs.

To generate the point source simulation, the biopsy image with the bone surface voxels labeled served as the base. A new three-dimensional voxelized image was then produced with the randomly chosen surface voxel labeled. Similarly, the evenly distributed source map was generated from the biopsy sample with all bone surface voxels labeled with the same value. The remaining two uneven distributions were built via a binomial distribution to give each voxel one of two values. The two scenarios chosen were 90% of the total activity randomized to 10% of the voxels, and 100% of the activity randomized throughout 50% of the voxels. This resulted in a total of four scenarios. To score the absorbed dose to each voxel within the samples, the GATE Dose Actor tool was used and attached to the biopsy.

### Simulation execution

Development of the model occurred over multiple testing and execution cycles to ensure repeatability of the simulations. Virtual Gate 9.3 and Geant4 version 11.1.1 was used for prototyping. A high-performance computing cluster with GATE 9.1, Geant4 10.7.1, and Root 6.20.02 (Antcheva et al. 2009) was employed to run the built simulations to allow for high volumes and quick simulation time.

### Analysis and interpretation

To visualize bone samples used in simulations, three-dimensional surface rendering of the micro-CT acquired bone biopsy were developed using the 3D Viewer volume plugin for FIJI (ImageJ 1.54f) (Schindelin, J., Arganda-Carreras, I., Frise, E. et al 2012). The volumes were thresholded to illustrate the bone surface structure. Samples were down resolved (2-3x binning).

Interpretation of absorbed doses are visualized through line profiles of total dose within concentric spheres surrounding a point of reference. One hundred spheres are plotted with radii set to 18 microns. Analysis is compared to bone structure visualized at the reference point with a representative slice, with Python 12.3 and matplotlib with the viridis colormap.

Three representative Radium-223 treated bmCRPC biopsy samples were selected. From these samples, we simulated four activity distributions and assessed the attendant dose profiles. The average activity per sample from our previous study (Benabdallah et al. 2024) was approximately 25 Bq, and this value was chosen in our experiments. We initiated simulations using the oversimplified case of assigning the complete sample activity to a single central voxel. As an opposite extreme scenario, we next investigated the dose distribution after equal distribution of the activity throughout the (interior) surface of the bone biopsy sample.

In order to simulate more realistic uptake patterns as may be expected within in vivo environments, we chose to utilize a binomial distribution to initialize activity as previous work demonstrates radium to accumulate irregularly at the bone-soft tissue interface due to physical and biological variables in the bone metastasis microenvironment. Two scenarios were simulated here: 90% of the total activity placed within 10% of the target region with the remaining 10% spread evenly throughout the bone tissue interface; and 100% of the total activity spread within 50% of the target surface area.

## Results

Visualization of the selected biopsy samples reveal a highly irregular structure, recapitulating the complex and highly structured bone architecture. Further heterogeneity is the result of diverse bone quality at these pathological sites of invading prostate adenocarcinoma (Figure 2).

Conversion of Hounsfield units to material in the GATE toolkit was completed via the Hounsfield material generator, which maps HU values to densities through a densities calibration curve and material data file (Supplemental Data). Further analysis on the biopsy samples revealed voxels of high HU values, above 1800. As these high values can impact simulation of transported particles an analysis was conducted, to quantify diiferences which may impact simulation results. Two 10 minute simulations with simplified cubical geometries were ran, one with all voxels set to 1000 and the second to 2000 and a total of 25 Bq was initialized at the center of the cube for each run. Both simulations exhibited low uncertainties at the point of activity placement (Supplemental Table 1), with the differences in absorbed dose and deposited energies reported to be small. The limited impact of these high HU voxels hence supports the continued analysis of the simulation results for these biopsy samples.

When the complete sample activity is assigned to a single voxel, we note that ablative absorbed doses can be observed, even in this short simulation run time (Figure 2). It is demonstrated that a maximum localized dose reaching up to 200 Gy for sample 1 and 300 Gy for samples 2 and 3. A sharp decrease is observed directly surrounding the source point, as would be expected from the short alpha particle pathlength. By approximately 60 microns, additional absorbed dose accrued is negligible.

Bone seeking radiopharmaceuticals unevenly adsorb to the bone surface. This is because the active surface area, consisting of bone formation and removal processes, is only a fraction of the total of the periosteum. As expected, the absorbed dose profile resulting from the homogeneous distribution of activity is more widely distributed than in the scenario of the total activity in a single voxel (Figures 3-5A). Simulations of heterogenous activity uptake resulted in observable differences in absorbed dose profiles as depicted in Figures 3-5C. For all three samples, structural elements visualized in Figures 3-5A, as well as the concentration of the activity appear to drive where dose is localized. Greater locally absorbed doses are measured from more concentrated regions as in the 10% scenario in comparison to the 50% scenarios.

Results of simulations from bone sample 1 are plotted and compared to the bone structure (Figure 3D). The evenly spread distribution graph illustrates the dose (Gy) distribution with a pronounced peak at the origin, sharply declining as the radius (mm) increases. The trend stabilizes to a near-horizontal line beyond 0.5 mm, indicating minimal dose distribution at greater distances. Similarly, the distribution consisting of 90% of activity in 10% of the source region shows the dose peaking sharply at the center but at a lower maximum value. The decline is rapid, although slightly more varied, suggesting concentrated activity in a smaller region. The third plot depicts the results of the simulation for 100% of the activity spread in 50% of the bone surface source region. This line exhibits the highest peak dose at the origin, with a swift drop-off. This indicates a broader spread of activity compared to the previous distribution graphs, but still concentrated. While all graphs show a steep initial decline, differences in peak height and decline sharpness reflect varying activity concentration and spread.

**Figure 2.**
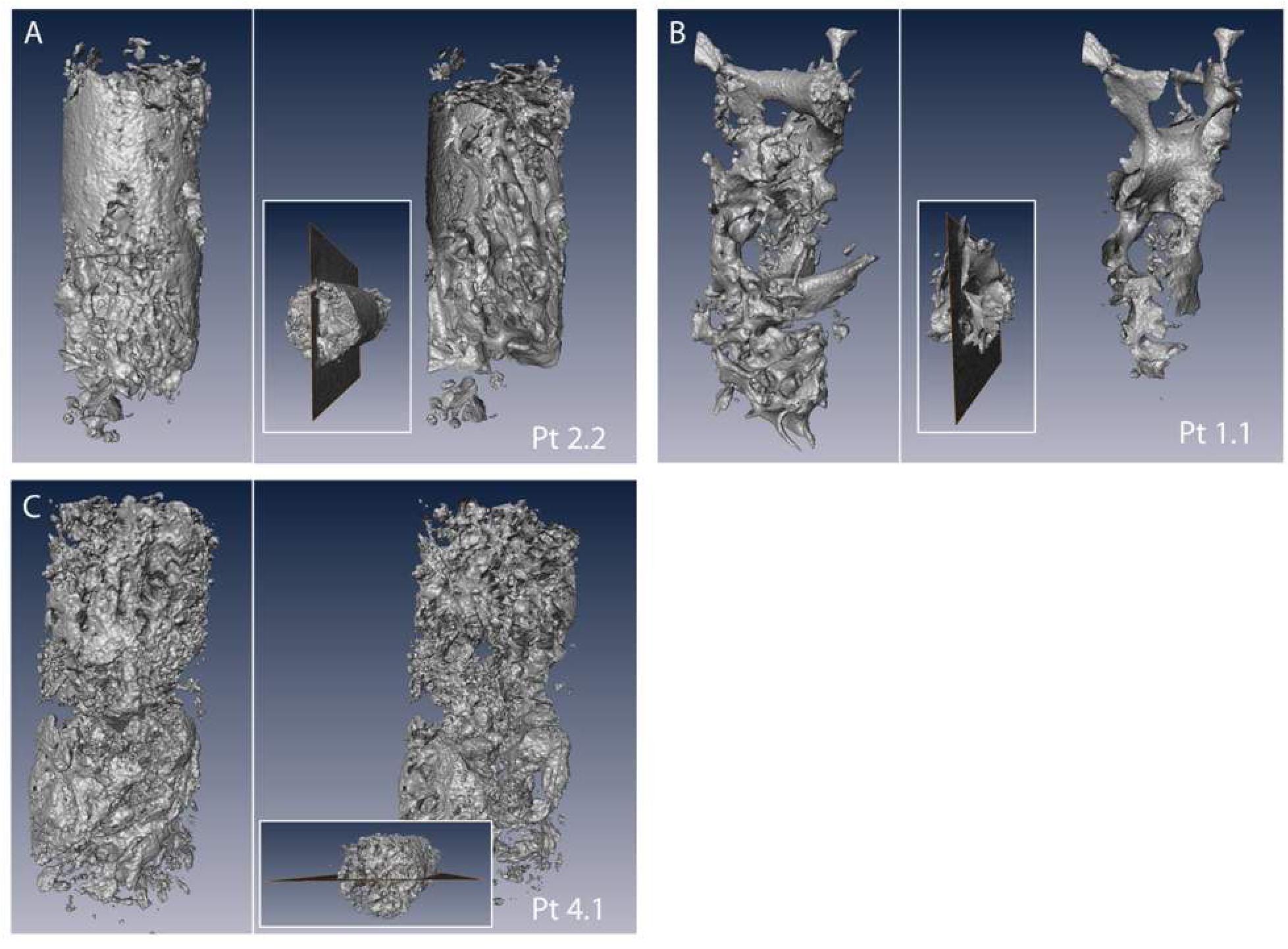
Biopsy Samples Visualized as Surface Projections. Two views of the bmCRPC bone biopsy are shown: rendered as an exterior surface, and as cut through the midline of the sample. Bone structure is noted to be highly irregular within, and differ significantly between, the samples.

Absorbed Dose surrounding point source for three simulations

**Figure 3.**
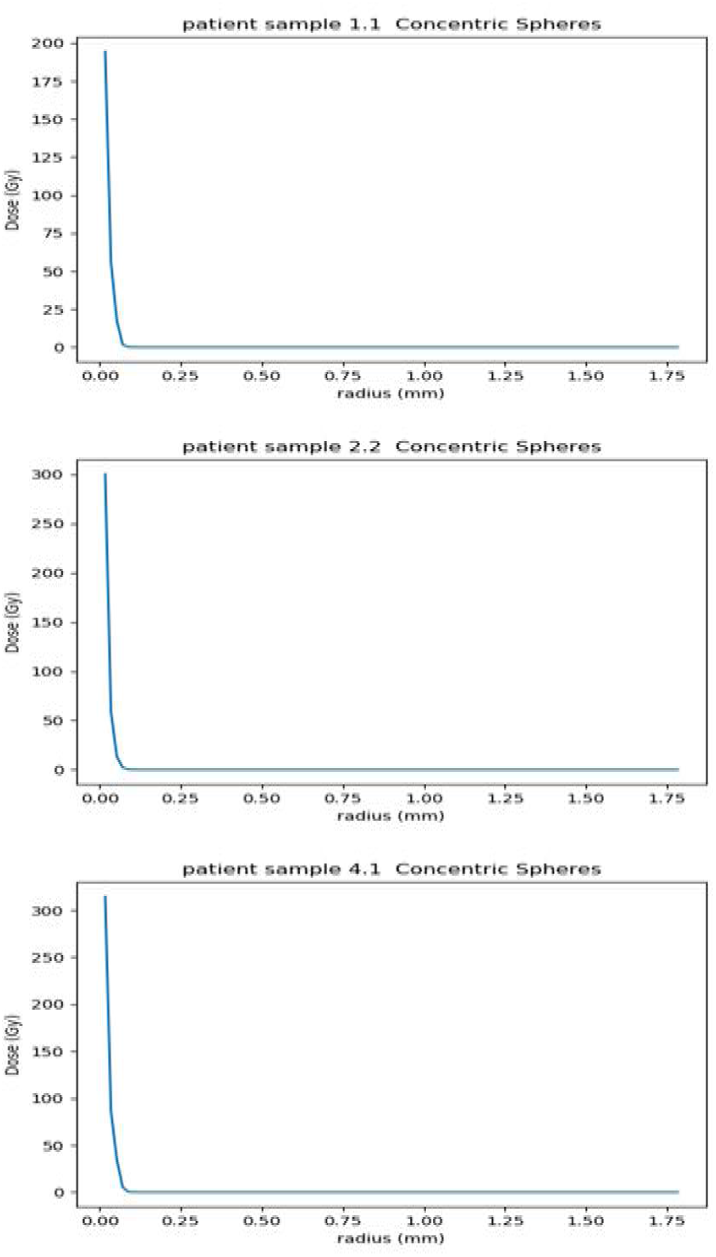
The absorbed dose is plotted within concentric spheres surrounding the point source. Similar patterns are noted for all three samples. Significant dose is measured at the point of activity with negligible amounts within 60 microns.

Absorbed dose for Patient Sample 1 in radiating spheres surrounding isocenter

**Figure 4.**
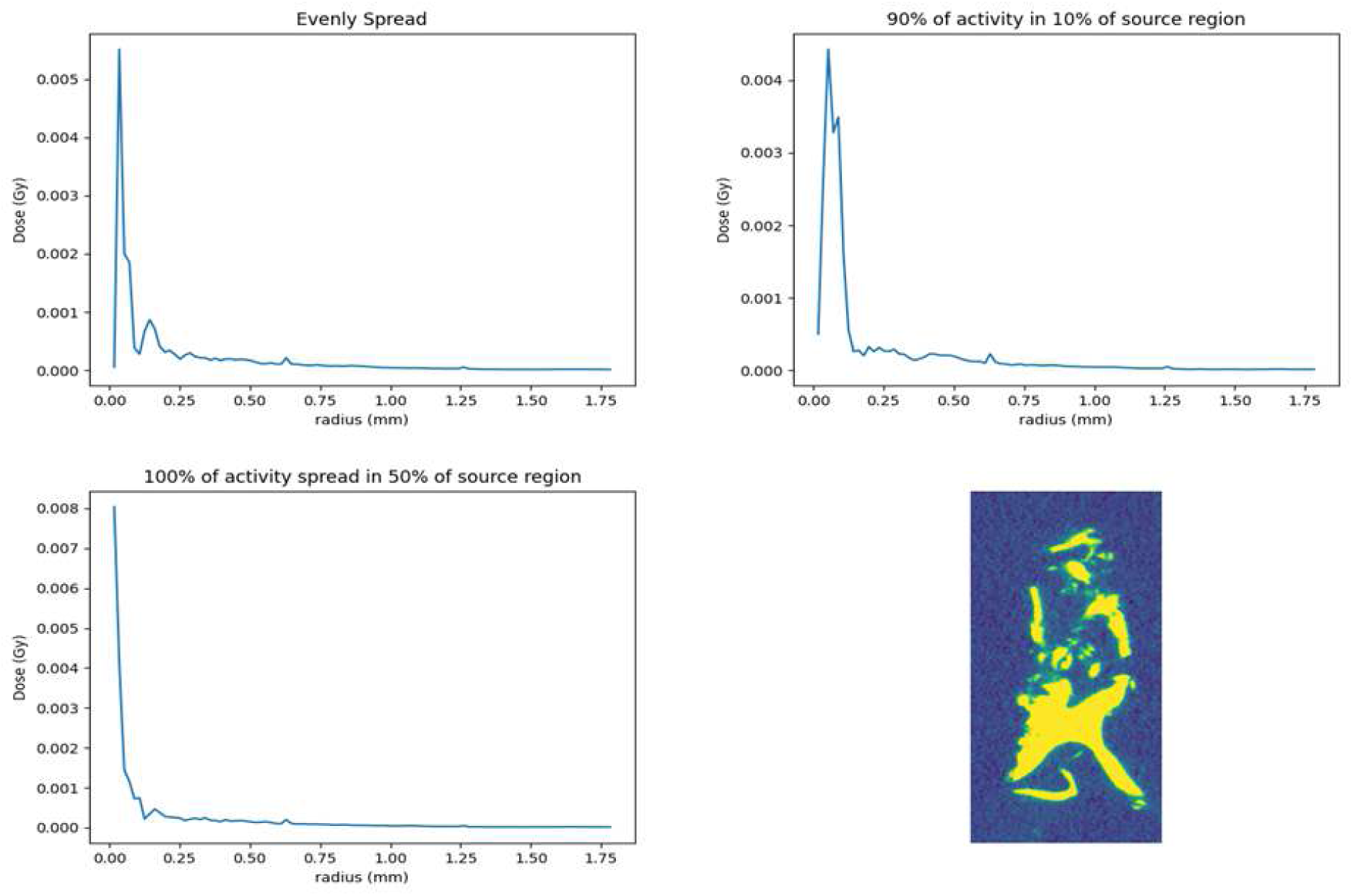
Dose profiles for three activity distributions are plotted for sample 1. The dose within concentric spheres surrounding the center of the sample is plotted as a function of radius. These three plots are compared to a slice of the bone at the isocenter.

Absorbed dose for Patient Sample 2 in radiating spheres surrounding isocenter

**Figure 5.**
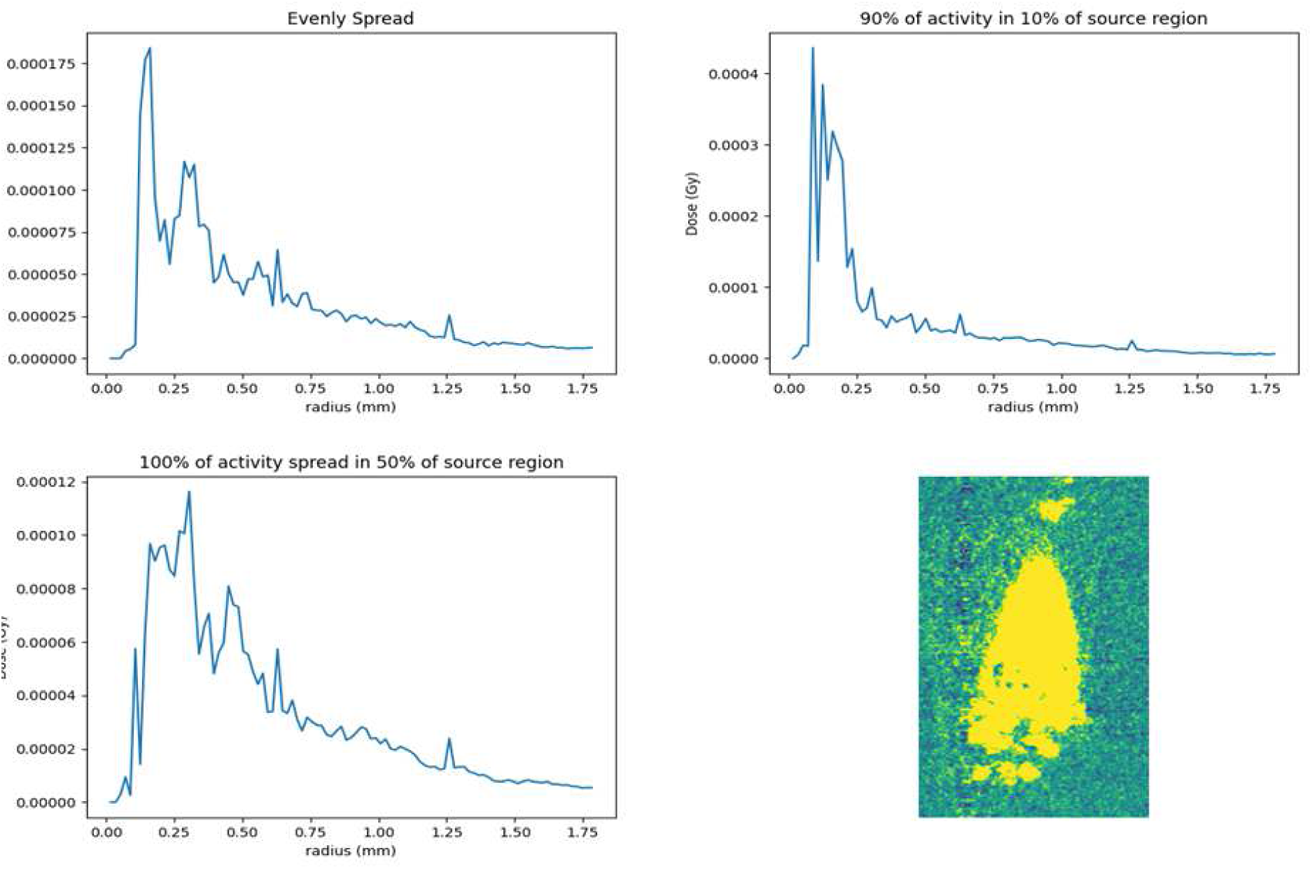
Sample 2 is depicted with the simulation outputs for three source distributions. Each line plot displays a dose distribution across different regions. Dose (Gy) is plotted against radius (mm).

Absorbed dose for Patient Sample 3 in radiating spheres surrounding isocenter

**Figure 5.**
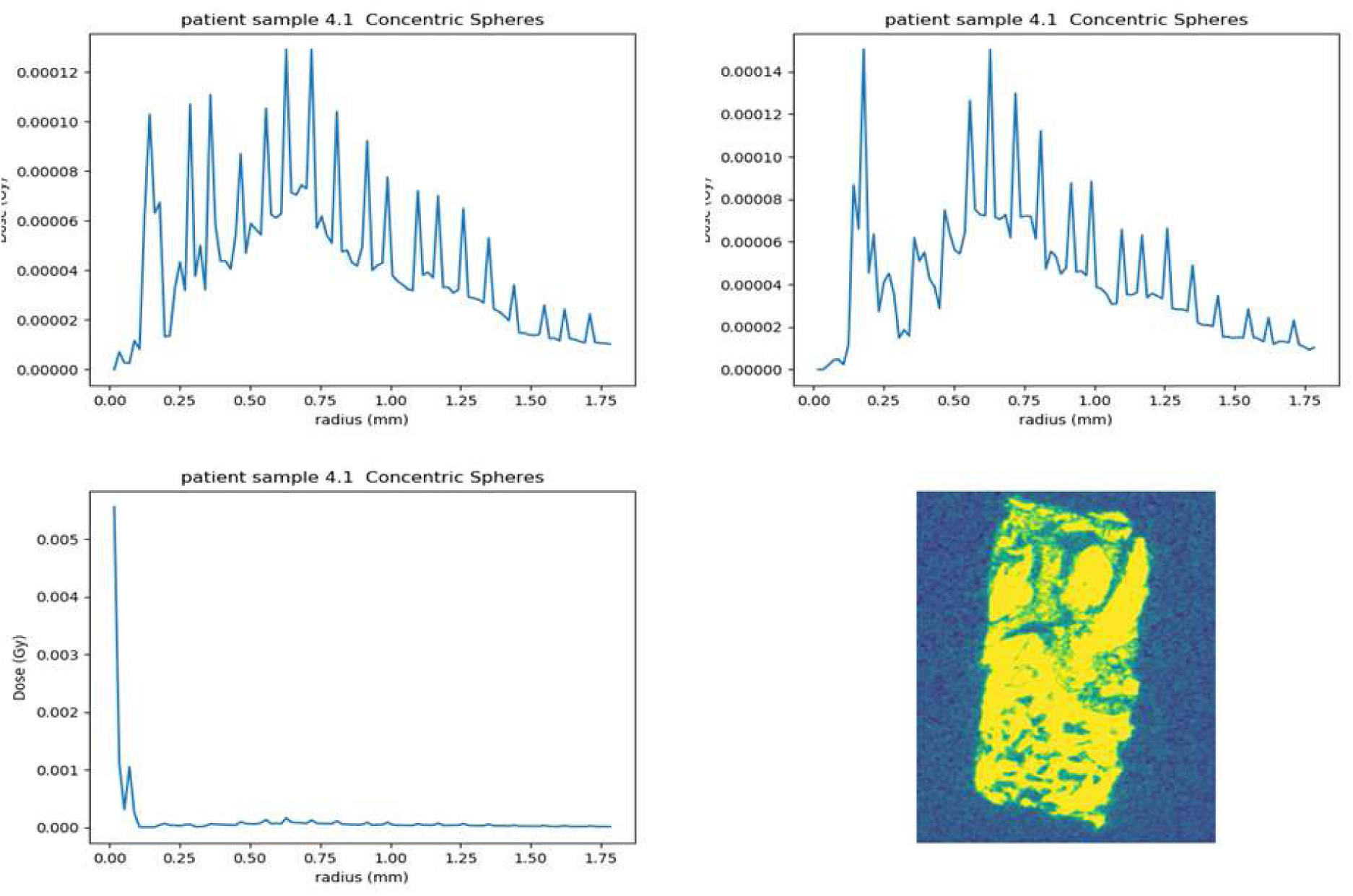
This representative patient samples displays different spread patterns and peak intensities of dose distributions across a specified radius than that observed with the other patient biopsy. Three plots are presented comparing radial dose distributions as well as a visualization of the bone at the isocenter.

Similarly, analysis of bone samples 2 and 3 recapitulate the results on differing samples. The mean and maximum dose values as well as the total deposited energy for the three tissue regions, (bone, bone surface, and soft tissue) for each of the three biopsy samples are found in Tables 1-4. The activity distributions in the four tables are labeled with values 1, 2, 3, and 4 using the following convention:

A. Point source
B. Homogenous spread
C. Heterogeneous spread: 90% of activity in 10% of the target region
D. Heterogeneous spread: 100% of activity in 50% of the target

**Table 1.**
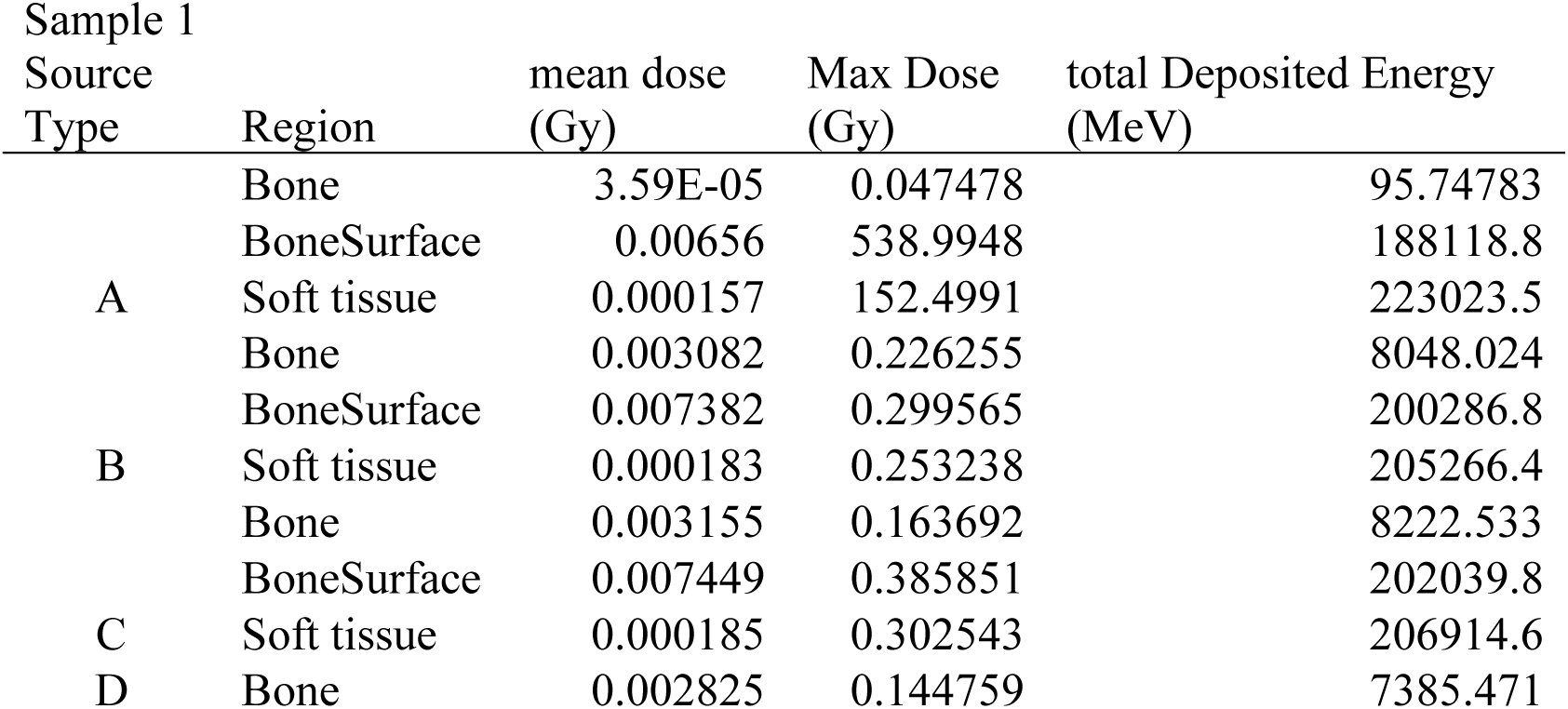
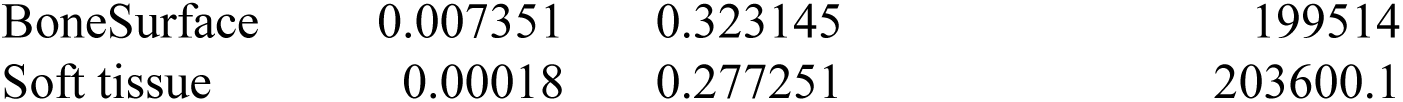
Sample 1 measurements for dose and deposited energy from four distribution patterns across the differing bone regions.

**Table 2.** Sample 2 measurements for dose and deposited energy from four distribution patterns across the differing bone regions.

| Sample 2 |  |  |  |  |
| --- | --- | --- | --- | --- |
| Source |  | mean dose | Max Dose | total Deposited Energy |
| Type | Region | (Gy) | (Gy) | (MeV) |
| A | Bone | 0.000285 | 129.3351 | 145110 |
|  | BoneSurface | 0.000729 | 1061.759 | 199063.4 |
|  | Soft tissue | 6.17E-05 | 137.253 | 37384.75 |
| B | Bone | 0.000197 | 0.192469 | 99370.45 |
|  | BoneSurface | 0.000929 | 0.221783 | 245583.2 |
|  | Soft tissue | 0.000114 | 0.190725 | 63462.49 |
| C | Bone | 0.000196 | 0.212065 | 98569.52 |
|  | BoneSurface | 0.00092 | 0.263121 | 243311.5 |
|  | NotBone | 0.000115 | 0.195434 | 64035.87 |
| D | Bone | 0.000195 | 0.1861 | 97925.34 |
|  | BoneSurface | 0.000925 | 0.276111 | 244456.1 |
|  | Soft tissue | 0.000116 | 0.229062 | 64487.55 |

**Table 3.** Sample 3 measurements for dose and deposited energy from four distribution patterns across the differing bone regions.

| Sample 3 |  |  |  |  |
| --- | --- | --- | --- | --- |
| Source Type | Region | mean dose (Gy) | Max Dose (Gy) | total Deposited Energy (MeV) |
| A | Bone | 0.001909 | 129.1936 | 59757.5 |
|  | BoneSurface | 0.004736 | 762.5891 | 233265.3 |
|  | Soft tissue | 9.96E-05 | 167.9435 | 120201.7 |
| B | Bone | 0.001145 | 0.365428 | 36023.87 |
|  | BoneSurface | 0.004558 | 0.370471 | 220717.8 |
|  | Soft tissue | 0.000114 | 0.348256 | 157397.2 |
| C | Bone | 0.001147 | 0.227777 | 36076.39 |
|  | BoneSurface | 0.004601 | 0.44939 | 222688.7 |
|  | Soft tissue | 0.000117 | 0.339412 | 159944.3 |
| D | Bone | 0.001155 | 0.278222 | 36270.1 |
|  | BoneSurface | 0.00455 | 0.368743 | 220355.1 |
|  | Soft tissue | 0.000114 | 0.332299 | 157284.3 |

**Table 4.** Sample 1 measurements for dose and deposited energy from four distribution patterns across the differing bone regions for a 2.4 hour simulation time.

| Sample 1 |  |  |  |  |
| --- | --- | --- | --- | --- |
| Source |  |  |  |  |
| Type | Region | Mean dose (Gy) | Max dose (Gy) | Total Deposited Energy (MeV) |
| A | Bone | 0.00055 | 1.704847 | 1473.112 |
|  | BoneSurface | 0.095077 | 7822.728 | 2726335 |
|  | Soft tissue | 0.002283 | 2174.153 | 3233952 |
| B | Bone | 0.042761 | 0.475256 | 111575.7 |
|  | BoneSurface | 0.106406 | 0.745203 | 2887510 |
|  | Soft tissue | 0.002625 | 0.566312 | 2946967 |
| C | Bone | 0.042262 | 0.611181 | 110248.2 |
|  | BoneSurface | 0.10608 | 1.341117 | 2878378 |
|  | Soft tissue | 0.002617 | 0.782224 | 2940296 |
| D | Bone | 0.042876 | 0.460481 | 111994.3 |
|  | BoneSurface | 0.10677 | 0.812104 | 2897884 |
|  | Soft tissue | 0.002627 | 0.640695 | 2953460 |

Here, the bone surface consists primarily of the bone tissue interface while voxels designated as not bone are predominantly mixed soft tissue at the lesion site. It is observed that when the dose and energy profiles are discriminated according to tissue type, greater values are consistently measured in the bone tissue interface. For all three samples, the greatest max dose is found for the point source while the scenario with homogenous and heterogenous spread results in the greatest mean values.

Table 4 displays similar trends for sample 1 for a 2.4-hour simulation; however, values are greater by an order of magnitude (Table 4).

Differences between the four distribution types (Point, Homogenous, Heterogenous: 90% activity in 10% target, and Heterogenous: 100% activity in 50% target) reveal several consistent trends across all three samples. For the Point source, the maximum dose and total deposited energy are significantly higher in the bone surface region compared to Bone and not bone regions. For the Homogenous distribution, the mean dose values are generally higher compared to the Point source, but the maximum dose and total deposited energy are lower. This suggests that a homogenous distribution spreads the radiation more evenly across the target regions, reducing peak doses but increasing overall exposure. The bone surface region still shows higher values compared to Bone and not bone, but the differences are less pronounced than in the Point source. In the Heterogenous distributions (both 90% activity in 10% target and 100% activity in 50% target), the trends are similar to the Homogenous distribution, with the BoneSurface region consistently showing higher values for mean dose, maximum dose, and total deposited energy. However, the heterogenous distributions exhibit slightly higher maximum doses and total deposited energy compared to the homogenous distribution, indicating that concentrating activity in specific areas increases localized radiation exposure. These consistent trends across all three samples validate the simulation tool in modeling a variety of activity uptake senereos in patient samples. Additionally, this further highlights the importance of distribution type in determining radiation dose and energy deposition patterns, which is crucial for optimizing radiation therapy and minimizing potential side effects.

## Discussion and Conclusions

Alpha particle emitting therapies are an active area of chemical and preclinical research, pharmaceutical development, and are of great clinical interest. Despite one agent, radium-223 dichloride citrate, having received approval from the EMA/FDA and many others in Phase I-III trials, there is limited information on the dose profiles within targeted cancer tissues or in surrounding healthy organs. This is important information that is expected to help guide the administered activities, lead to improved delivery approaches, and enable comparison with conventional therapeutic modalities.

Current approaches to define the absorbed dose in organs at risk and within lesions principally assume homogeneous distributions of activity. Such traditional methods to estimate absorbed dose for treatment effect or toxicity via whole organ analysis exclude the microdosimetric heterogeneity arising from the short pathlength and high linear energy transfer of alpha particles, as well as their inhomogeneous distribution at the cell scale. Using collected information on both the structure of the bone at sites of bmCRPC lesions, and the activity within these lesions, we have simulated dose profiles predicated on activity assignments under various scenarios. The result is a novel tool for preclinical development of alpha particle radio pharmaceuticals.

This work has demonstrated a novel preclinical approach for structural physics-based simulations. Via deployment of Geant4 software package and GATE, we successfully modeled Radium-223 dichloride treatment on patient bone samples. This work has also confirmed the importance of considering microstructural scale variabilities in alpha therapy development. In the aforementioned simulations with heterogeneous activity distributions, resultant dose maps differ from cases with activity spread evenly through the sample. It can be noted that absorbed dose variabilities throughout the samples are highly influenced by differences in structure of the bone. Anatomical differences at the micron level are known to vary significantly between individuals and especially in the microenvironment of disease sites.

These results underline the importance of accounting for microstructure in preclinical development of targeted radiotherapies. This is a difficult task, where the bone tissues sampled in the context of this study provided a unique means of contrast to assess structure via X-ray attenuation. Additionally, the absorption of delivered dose in the target region is an important factor to consider. In these simulations, we observe in scenarios with significant source variabilities, namely the scenario with half of all target voxels with no uptake, the localized absorbed doses are less concentrated. Further, confining all activity to a point source results in significant delivered dose near 300 Gy. The level of uptake of radium activity within the sampled biopsies and the low absorbed dose measures seen in simulations with activity spread throughout the sample suggests further analysis on increasing delivered activity may be advantageous to reach lethal levels of radiation, while for short durations. Expansion of this work will investigate additional biological targets and alpha emitting nuclide candidates.

## Acknowledgements

This work was funded in part by NIH NCI R01CA229893 and R01CA240711.

## Appendix A

### Material table

[Elements]

Hydrogen Carbon Nitrogen Oxygen Sodium Magnesium Phosphor Sulfur

Chlorine Argon Potassium Calcium

Titanium Copper Zinc Silver Tin

[/Elements]

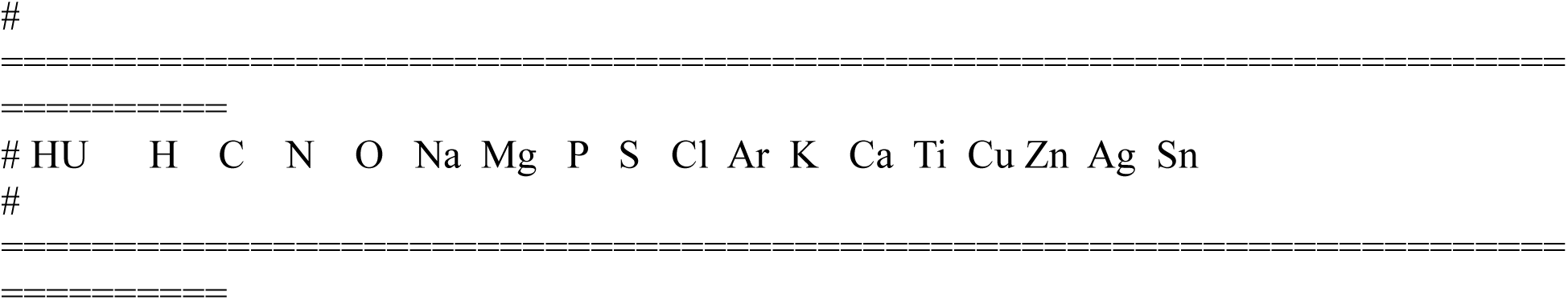
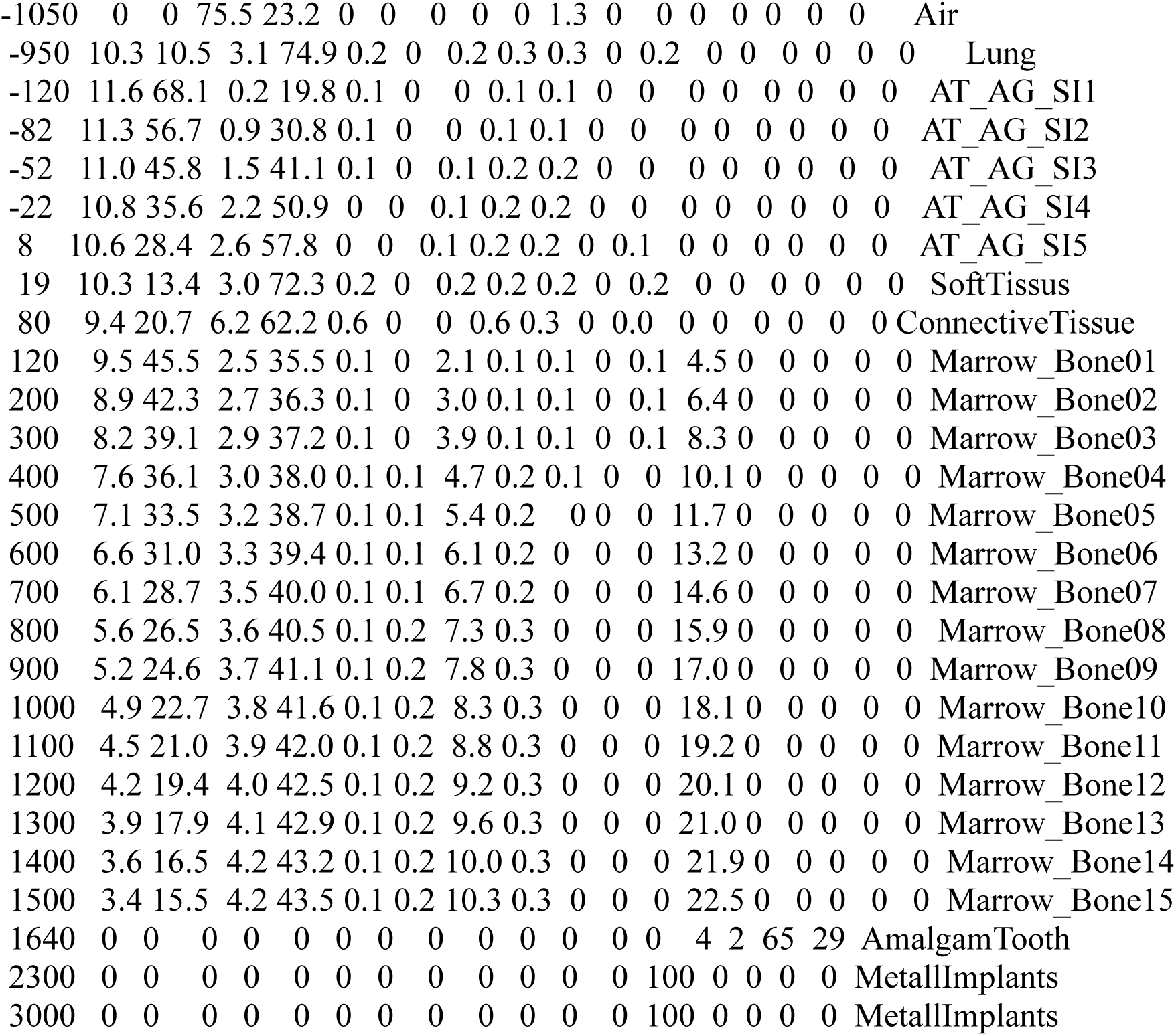

### Densities table

|  |  |  |
| --- | --- | --- |
| 503 | # | ===== |
| 504 | # HU | density g/cm3 |
| 505 | # | ===== |
| 506 | -1000 | 1.21e-3 |
| 507 | -98 | 0.93 |
| 508 | -97 | 0.930486 |
| 509 | 14 | 1.03 |
| 510 | 23 | 1.031 |
| 511 | 100 | 1.119900 |
| 512 | 101 | 1.076200 |
| 513 | 1600 | 1.964200 |
| 514 | 3000 | 2.8 |

**Supplemental Table 1:**

| HU value | Energy<br>(Mev) | Dose<br>(Gy) |
| --- | --- | --- |
| 1800 | 385654.3 | 0.030317 |
| 2500 | 392075.7 | 0.024808 |
| percent<br>difference | 1.665052 | -18.1701 |

